# Effects of VCD-induced ovarian failure on single muscle fiber contractility in a mouse model of menopause

**DOI:** 10.1101/2023.04.03.535419

**Authors:** Parastoo Mashouri, Jinan Saboune, W. Glen Pyle, Geoffrey A. Power

**Author notes:** Correspondence: Geoffrey A. Power PhD. Neuromechanical Performance Research Laboratory Department of Human Health and Nutritional Sciences College of Biological Sciences University of Guelph, Ontario, Canada.

## Abstract

**Objective:** Menopause is associated with impairments in muscle contractile function. The temporal and mechanistic basis of this dysfunction are not known. Using a mouse model of menopause we identified how gradual ovarian failure affects single muscle fiber contractility.

**Study design:** Mice were injected with VCD over 15 days and ovarian failure developed over 120 days. Mice were then sacrificed and slow-type soleus (SOL) and fast-type extensor digitorum longus (EDL) muscles were dissected and chemically permeabilized for mechanical testing.

**Main outcome measures:** Muscle fiber contractility was assessed via: force, rate of force redevelopment, instantaneous stiffness, and calcium sensitivity across three relative force levels (pCa_10_,pCa_50_,pCa_90_).

**Results:** Peak force and cross-sectional area (CSA) of the SOL were ∼33% and ∼24% greater in the VCD group as compared with controls (P<0.05), respectively, with no differences in force produced by the EDL fibers across groups (P>0.05). Upon normalizing force to CSA there were no differences across groups (P>0.05). Rate of force development was ∼33% faster for SOL in the VCD group compared to control. Ca^2+^ sensitivity did not differ between groups for either muscle at pCa_50_ (P>0.05). In the VCD group, Ca^2+^ sensitivity was higher for EDL, but lower for SOL at pCa_10_ and pCa_90_ (P<0.05), respectively.

**Conclusions:** In our mouse model of menopause, alterations to muscle contractility were much less evident as compared with ovariectomized models. This divergence across models highlights the importance of better approximating the natural trajectory of menopause during and after the transitional phase of ovarian failure on neuromuscular function.

## 1. Introduction

Menopause is the halt of menstruation associated with a decline in ovarian follicular activity, namely a decrease in the production of 17β-estradiol[1], which can have negative effects on muscle function[2]. Ovariectomized (OVX) rodent models are commonly used to investigate the effects of menopause on muscle contractility, however this approach does not approximate the natural trajectory of menopause nor the complex hormonal changes that occur during and after the transitional phase of ovarian failure. In this study we reduced circulating ovarian hormones through a model of gradual ovarian failure, allowing for investigation of the effects of the perimenopausal transition on muscle contractile function.

17β-estradiol deficiency is generally associated with reductions in both absolute and normalized (specific) force production in skeletal muscle[2–6]. Using an OVX model of ovarian failure in rodents, reductions in absolute force by ∼10-20% for the soleus (SOL) and extensor digitorum longus (EDL) muscles have been reported[7–9]. However, others using similar techniques reported no difference in absolute force for the SOL[10, 11] or EDL[10] muscles. Interestingly, some have even reported absolute force values ∼20% higher for the EDL muscle in an OVX group as compared to controls[12]. Additionally, when force is normalized to muscle mass, ‘specific force’ for both SOL and EDL muscles can be 9-30% lower[7, 8, 10] than controls for both whole muscle[10] and single muscle fibers[7, 8], indicating impairments in intrinsic contractile processes. As well, there have been reports of no difference in specific force with reductions in circulating ovarian hormones[11, 13, 14]. In studies in which reduced circulating 17β-estradiol levels caused reductions in absolute and specific force, 17β-estradiol replacement protected against these impairments in force generation[2, 8, 10, 15–18].

Cross-bridge based mechanisms are suggested to contribute to impaired muscle force production in both OVX and VCD induced models of ovarian failure[18]. The reductions in circulating 17β-estradiol levels are associated with reductions in strongly bound cross-bridge states by ∼10-15%[7, 8, 19], resulting in less force per myosin-actin interaction. Additionally, the proportion of attached cross-bridges are reduced as indicated by decreased active stiffness[7, 8]. The reduction in whole muscle and single fiber active stiffness coupled with a lower proportion of strongly-bound cross-bridges may explain force loss in the absence of circulating ovarian hormones[4, 5, 7, 9, 20–23]. Meanwhile, myofibrillar calcium (Ca^2+^) sensitivity does not seem to contribute to force loss in the OVX model for the SOL muscle[9]. However, it is unknown whether fast type muscle is affected, or whether VCD induced ovarian failure affects muscle contractility differently than the OVX model. Recently, Peyton et al. 2022 [19] observed, using the OVX model, that reductions in 17β-estradiol led to changes in the skeletal muscle phosphoproteome, including the phosphorylation of sarcomeric proteins involved in phospho-signaling networks for force generation, especially Ca^2+^-sensitive proteins. Thus, menopause induced reductions in muscle force generation may, in part, be attributed to changes in Ca^2+^ sensitivity of the muscle[19].

Importantly, OVX does not mimic the prolonged and complex hormonal transition that includes a retention in androgen production by the ovaries, and typical investigations occur 4-8 weeks after OVX, representing a chronic model of estrogen-deficiency. Our VCD model allows for an earlier investigation in which the effects of the perimenopausal transition can be examined. This study is the first investigation of this early and more physiologically relevant phase of menopause, in which we characterize the effects of the perimenopausal transition on single muscle fiber contractile function from a slow-type muscle (SOL) and fast-type muscle (EDL) using a VCD-induced ovarian failure mouse model.

## 2. Materials and Methods

### 2.1. Animals

Sexually mature CD1 female mice aged 78-105 days were obtained from Charles River Laboratories (St. Constant, QC), and housed on a 12-h light/dark cycle. Food and water were provided ad libitum. All procedures conducted were in accordance with the guidelines set by the Animal Care and Use Committee of the University of Guelph (AUP:4714) and the Canadian Council on Animal Care.

### 2.2. 4-Vinylcyclohexene diepoxide (VCD) mouse model of menopause

Mice were weighed and given daily intraperitoneal injections (160mg/kg) of VCD (MilliporeSigma, Oakville, ON) for 15 consecutive days[24]. VCD accelerates follicular atresia by inducing the selective loss of primary and primordial follicles in the ovary but leaves intact the rest of the ovary for residual hormone production. VCD mediates its ovarian effects by inhibiting autophosphorylation of membrane receptor c-kit[25]. Vaginal cytology was used to confirm lack of estrus cycles indicating ovarian failure[26, 27]. Mice were considered acyclic after 10 consecutive days of persistent diestrus which was achieved by day 120 representing the end of perimenopause and the start of menopause[28]. Muscle samples from all VCD mice were taken at day 120 following the start of VCD injections.

### 2.3. Tissue Preparation

The soleus (SOL), and extensor digitorum longus (EDL), muscles were harvested following sacrifice and were placed in chilled dissecting solution, chemically permeabilized and stored as reported previously[29]. *Please see the supplemental data page for all solution recipes*.

### 2.4. Mechanical testing and force measurements

Single fibers were transferred into a temperature-controlled chamber (15°C) and tied between a force transducer (model 403A; Aurora Scientific) and a length controller (model 322C; Aurora Scientific). Average sarcomere length (SL) was measured using a high-speed camera (Aurora Scientific). Fiber length (L_0_) was recorded, and fiber diameter was measured at three different points along the fiber using a reticule on the microscope in relaxing solution to calculate cross-sectional area (CSA) assuming circularity. First, a ‘fitness’ contraction was performed at 2.5μm in pCa4.5, after which SL was re-measured and, if necessary, re-adjusted to optimal rodent hindlimb SL, 2.5μm. To initiate activation, the fibers were transferred to a pre-activating solution (reduced Ca^2+^ buffering capacity with ATP) for 20s, then to an activating solution (varying levels of Ca^2+^ and high ATP) for 30s[29]. The highest force reached during the 30s window was taken as optimal force (P_o_) at each pCa level.

To determine the force-pCa relationship, 7 pCa levels (7.0-4.5) were used in a random order. Data were fit using a modified Hill equation:

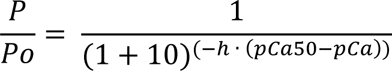

The pCa value at which 10%, 50% and 90% of maximal force was elicited (pCa_10_, pCa_50_, pCa_90_) was used to determine differences in Ca^2+^ sensitivity.

The rate of force redevelopment (k_tr_) was assessed using a slack re-stretch method[30, 31].

By rapidly shortening the fiber with a ramp of 10 L_o_/s by 15% of L_o_ and then a rapid (500 L_o_/s) re-stretch back to L_o_, we force all cross-bridges to break, and then the re-stretch allows further dissociation of any remaining cross-bridges. The force redevelopment is related directly to the reattachment of myosin to actin and a redistribution of cross bridges from pre-power stroke into force-generating states. A mono-exponential equation, y=a(1-e^-kt^)+b, was fit to the redevelopment curve to determine k_tr_.

To assess the proportion of attached cross-bridges, instantaneous stiffness (*k*) tests were performed by inducing a rapid (500 L_o_/s) stretch of 0.3% of L_o_ and dividing the change in force during the stretch by the length step. Instantaneous stiffness was calculated as the difference between peak force during the stretch and the average force over the 500ms prior to the stretch, divided by the change in fiber length induced by the stretch. Absolute force was reported as the average force over 500ms during the steady-state phase. All force measures were adjusted for resting passive tension by subtracting the average baseline value for the first 50ms of the trial in the relaxing solution, thus for all contractions, active force is reported. If at low [Ca^2+^] a negative value was recorded owing to any force drift of the transducer, the value was replaced with ‘0’. All force values were normalized to CSA for calculations of specific force.

If force decreased by >10% from the maximal force produced at the beginning of the experiment, or if the striation pattern of the muscle fibers became unclear such as to not allow measurements of sarcomere length, the experiment was terminated. Any fiber that slipped or ripped before the completion of the tests was removed from the testing apparatus and excluded. A total of N=47 fibers from the EDL and N=55 fibers from the SOL were included in the analysis.

### 2.5. Fiber Typing

Following mechanical testing, fiber typing was performed[29, 32, 33]. Gels were run at a constant voltage of 50V for approximately 40h at 4°C. Fibers containing MHC I and MHC II were classified as Type I and Type II, and were determined by comparing to a standard protein ladder (Bio-Rad Protein Plus Standard 10-250 kD) with known molecular weights.

### 2.6. Statistical Analysis

2-way ANOVAS (PRISM 9.5.0) were run to assess differences between groups (VCD, control) and across muscles (SOL, EDL) for: absolute force, specific force, ktr, stiffness, fiber CSA, fiber types (within SOL), pCa_10_, pCa_50_, and pCa_90_ values. An alpha level of 0.05 was used.

## 3. Results

### Absolute force

There was an interaction of group×muscle (F(1,99)=6.482, p=0.012), and a main effect of group (F(1,99)=4.802, p=0.031) such that in the VCD group, fibers from the SOL, produced ∼33% more force than the control group (Figure-1A). As well, there was no main effect of muscle (F(1,99)=0.0575, p=0.811). For the SOL, when fiber type was taken into consideration, there was an interaction of fiber type×group (F(1,51)=4.875, p=0.032), and a main effect of group (F(1,51)=18.540, p<0.001), such that within the Type I fibers, the VCD group produced ∼71% force more than the control group. Additionally, within the VCD group, the Type I fibers produced ∼24% more force compared to Type II fibers (Figure-1B).

**Figure 1.**
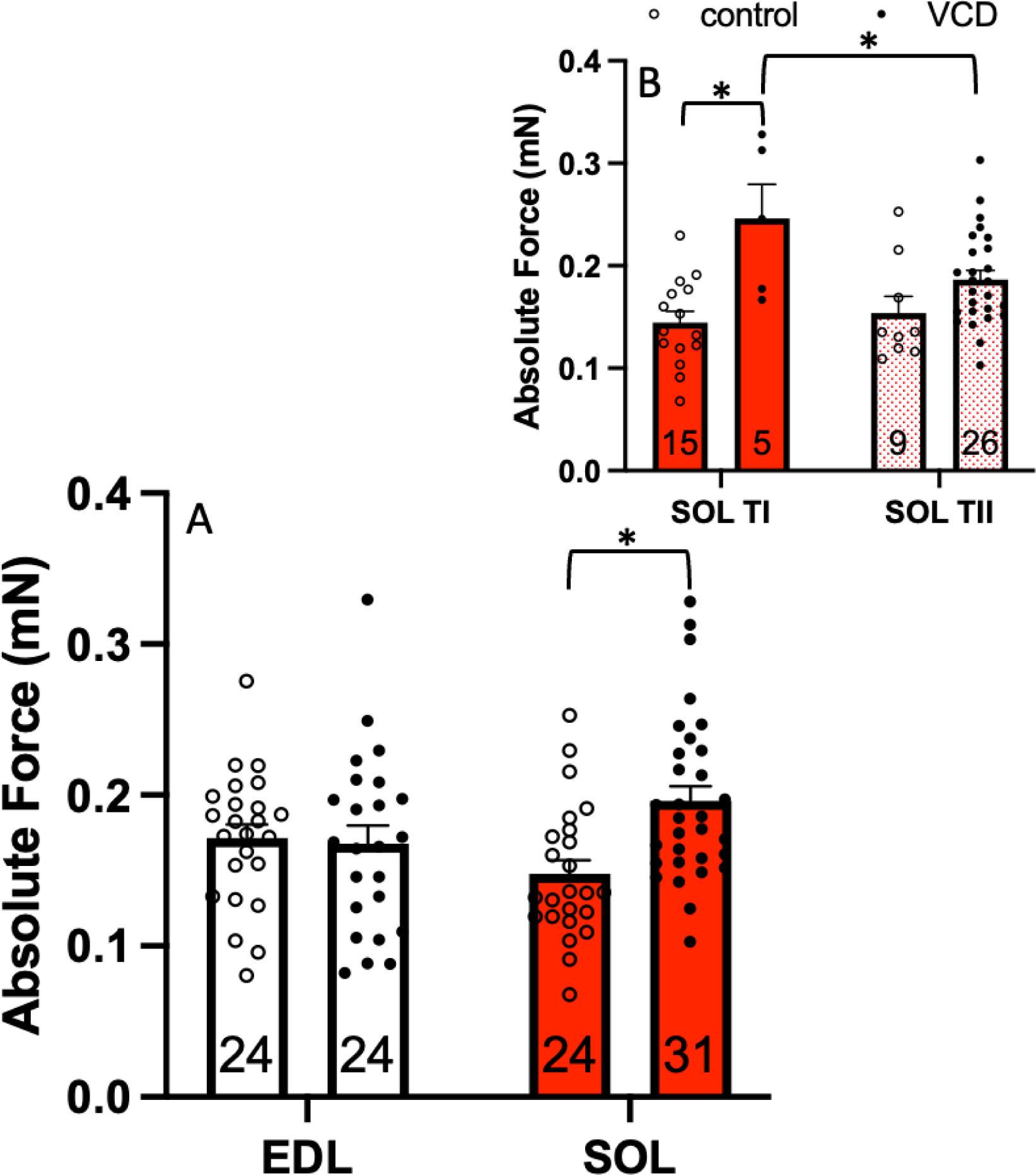
A) Absolute force: For the VCD group there was ∼33% greater maximal force production for fibers from the SOL muscle as compared with the controls. There were no differences for the EDL muscle across groups. EDL ctrl, n=24; EDL VCD, n=24; SOL ctrl, n=24; SOL VCD, n=31. B) SOL muscle: When analyzing the SOL muscle by fiber type, TI fibers produced ∼71% more force than the control group, and the TI fibers from the VCD group produced ∼24% more force than TII fibers from the VCD group. Type I ctrl, n=15; Type I VCD, n=5; Type II ctrl, n=9; Type II VCD, n=26. * Indicates significant difference (P<0.05).

### Cross-sectional area

There was an interaction of group×muscle (F(1,99)=5.048, p=0.027) (Figure-2A), with no main effects of muscle (F(1,99)=0.772, p=0.382) or group (F(1,99)=0.789, p=0.377). In the VCD group, fibers from the SOL were ∼24% larger than the control group (Figure-2A). For the SOL, when fiber type was taken into consideration, there was no interaction of fiber type×group (F(1,51)=0.246, p=0.662), main effects of group (F(1,51)=2.209, p=0.143) or muscle (F(1,51)=0.234, p=0.631) (Figure-2B).

**Figure 2.**
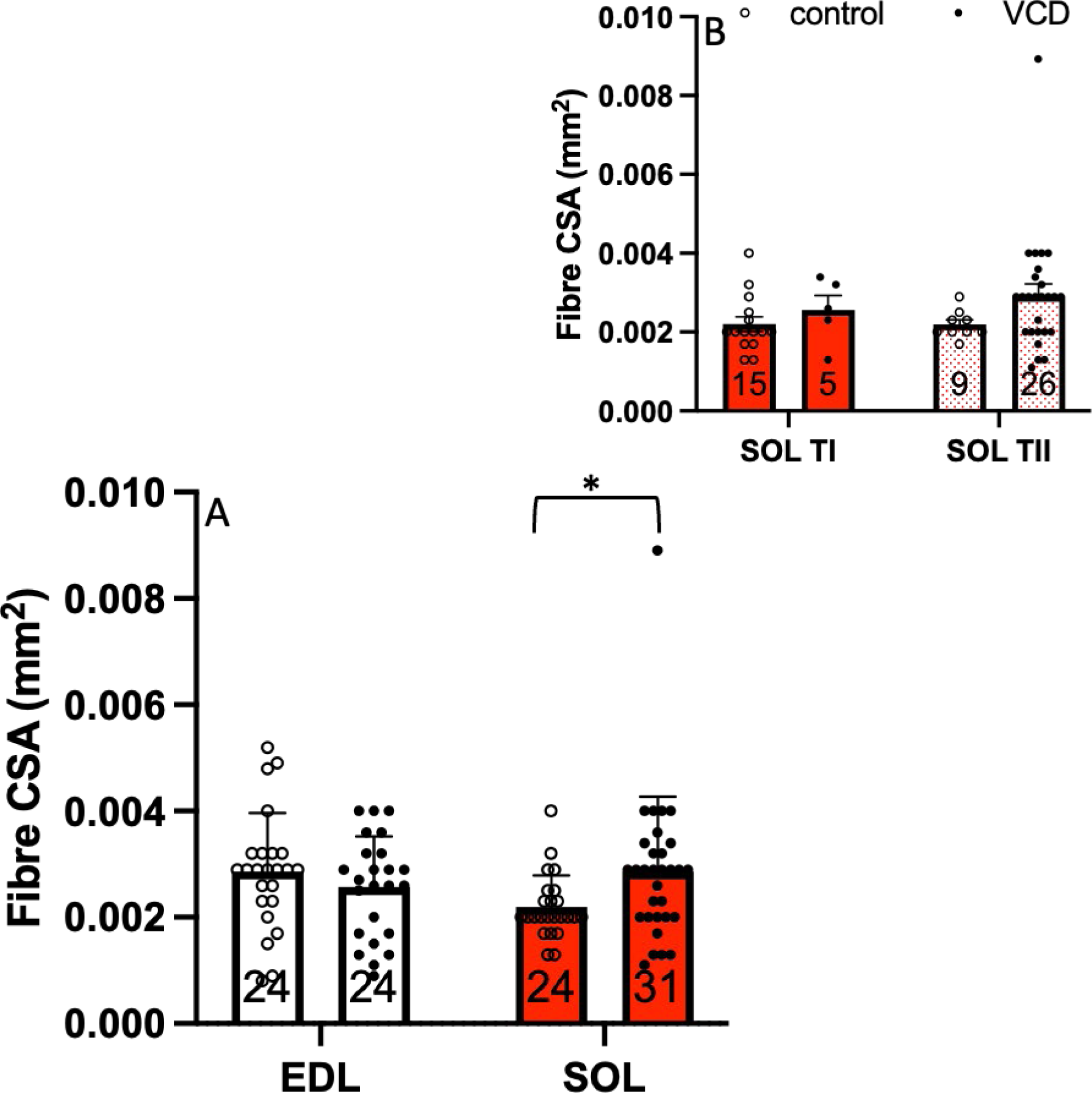
A) Fiber CSA: For the VCD group there was ∼24% larger CSA for fibers in the SOL muscle as compared with the controls. EDL ctrl, n=24; EDL VCD, n=24; SOL ctrl, n=24; SOL VCD, n=31. B) SOL muscle: When Fiber type was taken into consideration, no significant differences between any of the groups or fiber types were observed. Type I ctrl, n=15; Type I VCD, n=5; Type II ctrl, n=9; Type II VCD, n=26. * Indicates significant difference (P<0.05).

### Specific force

There was no interaction (F(1,99)=0.997, p=0.320) or main effects of group (F(1,99)=1.451, p=0.231) or muscle (F(1,99)=1.188, p=0.278) (Figure-3A). Therefore, when force was normalized to CSA to account for myofibrillar protein content, differences in absolute force were eliminated. For the SOL, when fiber type was taken into consideration, there was no interaction of fiber type×group (F(1,51) =2.956, p=0.092), or main effects of muscle (F(1,51)=1.149, p=0.289). However, there was a main effect of group (F(1,51)=5.012, p=0.030), indicating that fibers from the VCD group produced ∼20% more specific force compared to control groups (Figure 3B).

**Figure 3.**
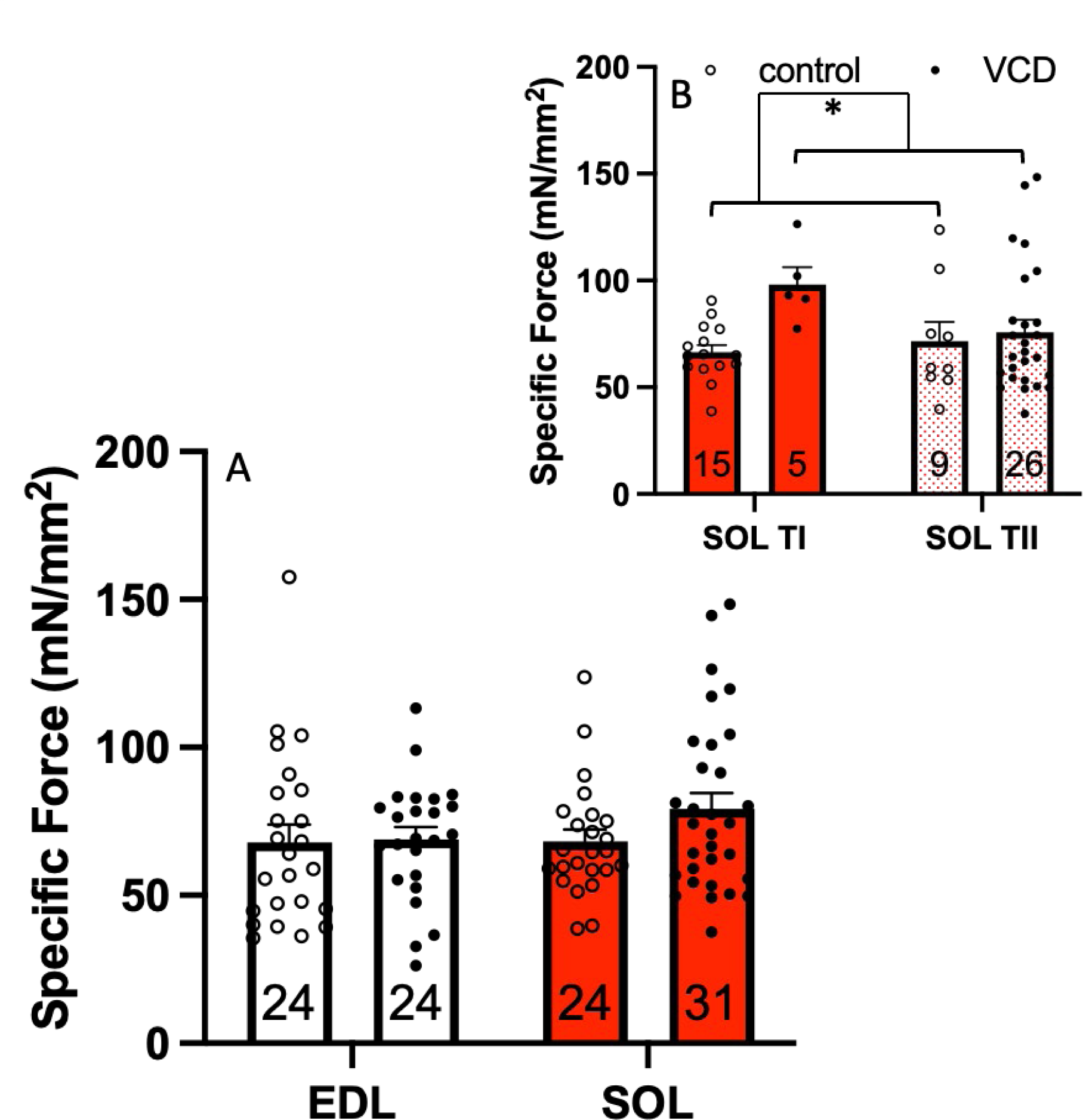
A) Specific force: When force was normalized to CSA to account for myofibular protein content, no differences in specific force were observed. EDL ctrl, n=24; EDL VCD, n=24; SOL ctrl, n=24; SOL VCD, n=31. B) SOL muscle: When fiber type was taken into consideration, the VCD groups produced ∼20% more force compared to controls. Type I ctrl, n=15; Type I VCD, n=5; Type II ctrl, n=9; Type II VCD, n=26. * Indicates significant difference (P<0.05).

### Instantaneous stiffness

There was no interaction of group×muscle (F(1,99)=0.0242, p=0.877) or main effect of group (F(1,99)=0.740, p=0.392). However, there was a main effect of muscle (F(1,99)=8.920, p=0.004), with fibers from the SOL having ∼20% greater instantaneous stiffness compared to fibers from the EDL muscle (Figure-4A). For the SOL, when fiber type was taken into consideration, there was no interaction of fiber type×group (F(1,51)=2.280, p=0.137), main effects of group (F(1,51)=1.428, p=0.238) or muscle (F(1,51)=1.058, p=0.309), suggesting no differences in the proportion of attached crossbridges within the SOL when comparing groups and fiber types (Figure-4B). However, coupled with the force data, this data likely suggests an increase in the proportion of strong-binding cross-bridges in the SOL compared to the EDL.

**Figure 4.**
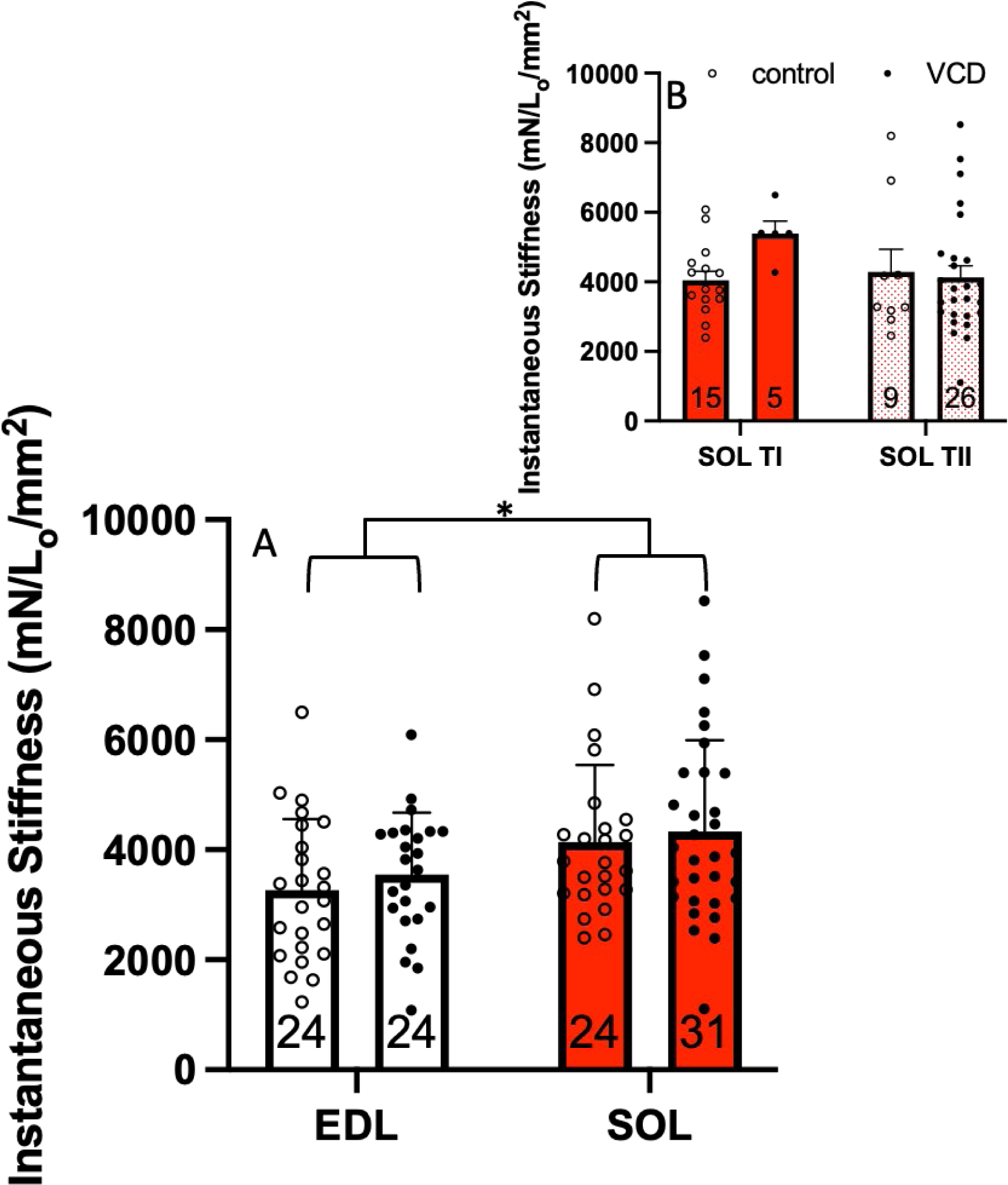
A) Instantaneous stiffness: The SOL muscle had ∼20% greater instantaneous stiffness compared to the EDL muscle, indicating a greater proportion of attached crossbridges. EDL ctrl, n=24; EDL VCD, n=24; SOL ctrl, n=24; SOL VCD, n=31. B) SOL muscle: When fiber type was taken into consideration, no significant differences were observed. Type I ctrl, n=15; Type I VCD, n=5; Type II ctrl, n=9; Type II VCD, n=26. * Indicates significant difference (P<0.05).

### Rate of force development

There was an interaction of group×muscle (F(1,99)=7.870, p=0.006), such that single fibers from the SOL had a 33% faster rate of force development in the VCD group as compared with the control group (Figure-5A). Additionally, there was the expected main effect of muscle (F(1,99)=58.60, p<0.001) such that fibers from the EDL produced on average ∼41% faster rate of force development as compared with SOL (Figure-5A). There was no main effect of group (F(1,99)=2.403, p=0.124). For the SOL, when fiber type was taken into consideration, there was an interaction of fiber type×group (F(1,51)=8.316, p=0.006), such that Type II fibers had ∼82% and ∼48% faster rate of force development compared to Type I fibers in the VCD and control groups, respectively (Figure-5B). Additionally, there was a main effect of fiber type (F(1,51)=70.32, p<0.001), such that Type II fibers produced ∼66% faster rate of force development as compared with TI fibers (Figure-5B).

**Figure 5.**
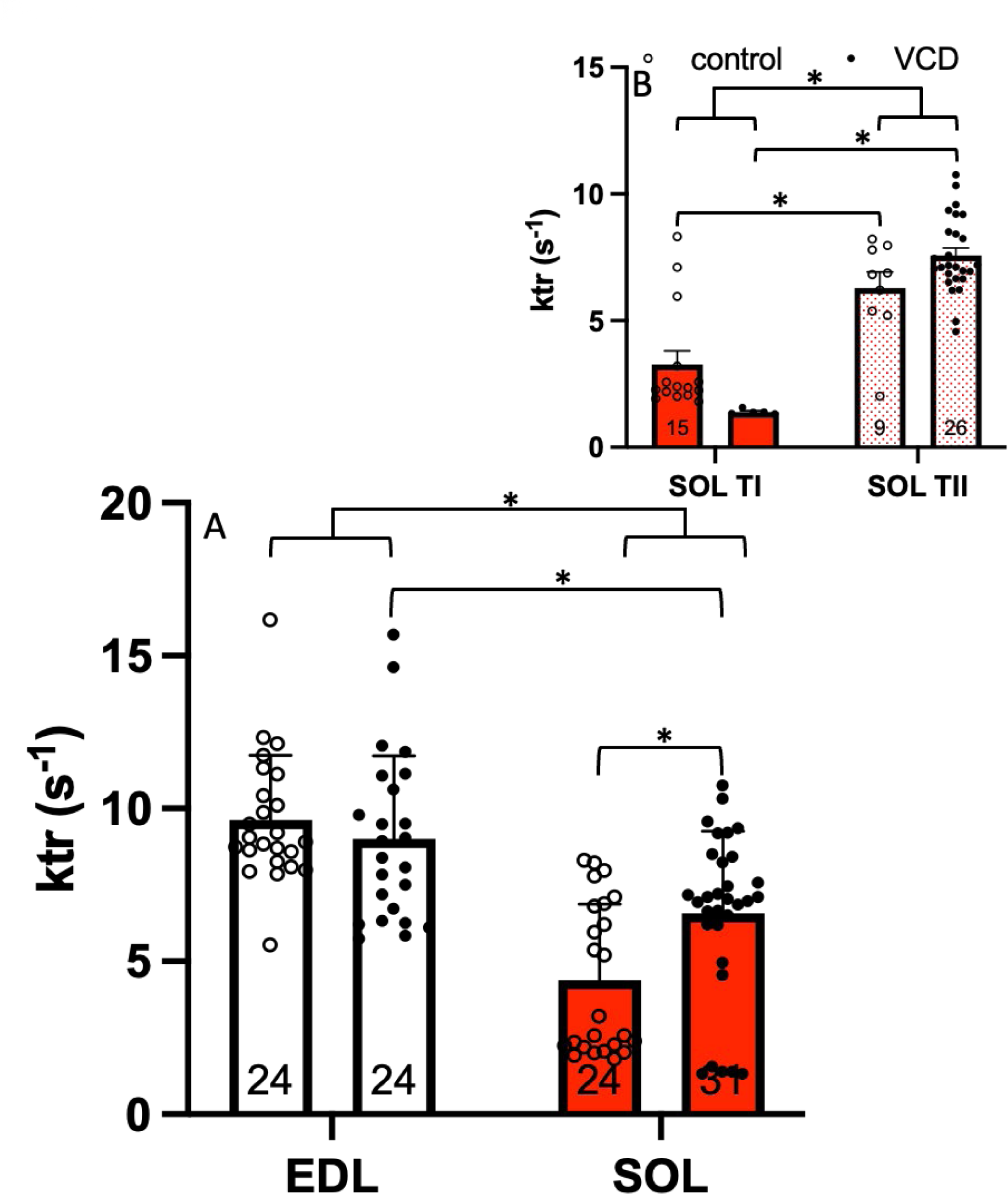
A) ktr: Single fibers from the SOL had a ∼33% faster rate of force development in the VCD group as compared with the controls. Additionally, fibers from the EDL muscle produced on average ∼41% faster rate of force development as compared with SOL. EDL ctrl, n=24; EDL VCD, n=24; SOL ctrl, n=24; SOL VCD, n=31. B) SOL muscle: When fiber type was taken into consideration, Type II fibers were ∼48% and ∼82% faster than Type I fibers in the control and VCD groups, respectively. Additionally, Type II fibers were ∼66% faster thab Type I fibers. Type I ctrl, n=15; Type I VCD, n=5; Type II ctrl, n=9; Type II VCD, n=26. * Indicates significant difference (P<0.05).

### Calcium sensitivity from low to high force output

For force production at low concentrations of Ca^2+^ (pCa_10_), there was an interaction of group×muscle (F(1,99)=5.801, p=0.018, with no main effects of group (F(1,99)=0.453, p=0.502), or muscle (F(1,99)=3.837, p=0.053) (Figure-6A). In the VCD group, the EDL was ∼5% more sensitive to Ca^2+^ compared to the SOL (Figure-6A). For force production at 50% of Ca^2+^ concentration (pCa_50_), there was no interaction of group×muscle (F(1,99)=0.0165, p=0.898), main effects of group (F(1,99)=1.863, p=0.175), or muscle (F(1,99)=3.209, p=0.076), indicating no change in Ca^2+^ sensitivity at a pCa concentration where 50% of maximum force is produced (Figure-6B). For force production at high Ca^2+^ concentrations (pCa_90_), there was no interaction of group×muscle (F(1,99)=2.319, p=0.131), or main effect of group (F(1,99)=2.247, p=0.137). However, there was a main effect of muscle (F(1,99)=23.78, p<0.001), whereby, in the VCD group the SOL was ∼6% more sensitive to Ca^2+^ compared to the EDL (Figure-6C). Thus, at pCa_90_, a Ca^2+^ concentration where 90% of maximum force is produced, the SOL was more sensitive to Ca^2+^ compared to the EDL.

**Figure 6.**
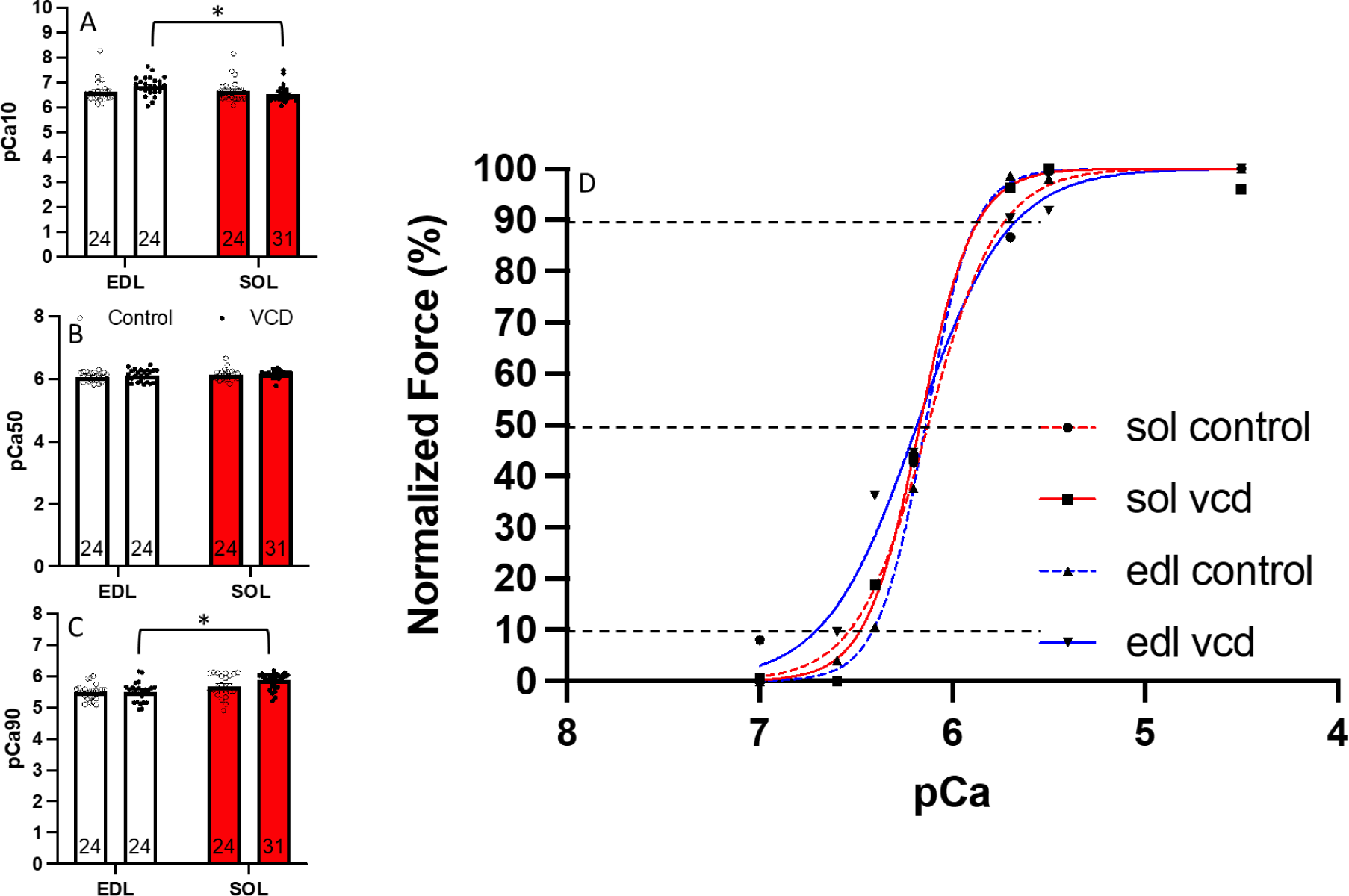
A) pCa10: In the VCD groups, the EDL muscle was ∼5% more sensitive to Ca^2+^ compared to the SOL muscle. EDL ctrl, n=24; EDL VCD, n=24; SOL ctrl, n=24; SOL VCD, n=31. B) pCa50: No significant differences were observed for the pCa50 value. EDL ctrl, n=24; EDL VCD, n=24; SOL ctrl, n=24; SOL VCD, n=31. C) pCa90: In the VCD groups, the EDL muscle was 6% more sensitive to Ca^2+^ compared to the SOL muscle. EDL ctrl, n=24; EDL VCD, n=24; SOL ctrl, n=24; SOL VCD, n=31. D) Representative Force-pCa curve displaying changes in force at varying pCa levels. pCa 4.5 demonstrates highest Ca^2+^ concentration, used for maximal activation of fibers. pCa 7.0 demonstrates lowest Ca^2+^ concentration. * Indicates significant difference (P<0.05).

## 4. Discussion

Using a VCD mouse model of ovarian failure to characterize the effects of the perimenopausal transition on single muscle fiber contractility, we found either an increase in contractile performance, or no change as compared with controls. In terms of muscle contractile function, this model of ovarian failure appears to yield divergent findings as compared with classic OVX post-menopausal (4-8 weeks post surgery) models[7, 8, 10, 11, 14, 18, 19], and is more in line with results from studies on human (49-57yrs) single muscle fibres [45, 46]. However, time-course changes are yet to be identified.

### 4.1. Ovarian failure led to an increase in force generation for the SOL

Reducing circulating ovarian hormones through the OVX model results in a reduction in absolute force production as compared with controls[7, 8, 10, 11, 14, 18, 19] on average 4-8 weeks following surgery. However, using a VCD-induced model of ovarian failure, we found the opposite during the menopausal transition, an ∼33% increase in SOL single muscle fiber force as compared with controls, and no change in force generation for fibers from the EDL. The increase in absolute force in the VCD group for the SOL may in part be due to the intact ovaries maintaining theca and granulosa cells[18, 34], which are responsible for producing testosterone, as the levels of ovarian estrogens declined. Thus, the relative proportion of testosterone to 17β-estradiol in the cells increase[34–39], likely leading to the larger CSA and greater absolute forces which we observed for the SOL. However, we did not observe a difference in specific force (force normalized to CSA) or instantaneous stiffness, indicating neither intrinsic contractile properties or the proportion of attached cross-bridges differed between groups. Impairments in force production using OVX models appear to be driven by a reduction in the total number of bound crossbridges, and less force per bridge [2]. However, these cross-bridge based alterations were not evident in the present study.

### 4.2. Calcium sensitivity is altered in a fiber-type dependent manner following ovarian failure

Reducing circulating ovarian hormones through the OVX model had no effect on Ca^2+^ sensitivity of SOL single muscle fibres 10-14 weeks after surgery as compared with controls[9]. However, it has been shown previously that Ca^2+^ sensitivity is lower in cardiomyocytes with 17β-estradiol deficiency[19, 40], and that 17β-estradiol deficiency generally leads to reductions in the phosphorylation of myosin regulatory light chain (RLC) [13, 19], likely affecting Ca^2+^ sensitivity in skeletal muscle[41]. In the present study, in the VCD group, fibers from the EDL were in fact more sensitive to Ca^2+^ at low force levels (pCa_10_), compared to the control group, and fibers from the SOL were more sensitive to Ca^2+^ at high force levels (pCa_90_) (Figure-6). The increase in Ca^2+^sensitivity in our study as compared to previous findings requires further investigation and may be related to changes in RLC phosphorylation.

### 4.3. Rate of force redevelopment increases following ovarian failure in the SOL muscle

It has been reported previously, that Ca^2+^ concentration affects k_tr_, such that as Ca^2+^ increases, so does k_tr_ until a plateau is reached[30, 32, 42]. In the current study, we found that for the SOL in the VCD group there was an increase in maximal rate of force development compared to the control group (Figure-5A). When fibers from the SOL were separated by fiber type, the control group had the expected ∼63% TI and 37% TII fiber type distribution. However, the fiber distribution for the SOL in the VCD group was 84% TII fibers (Figure-7). Therefore, it is likely that the reduction in 17β-estradiol caused a shift in the proportion of fiber types, driving the increase in rate of force development in the SOL of the VCD group.

**Figure 7.**
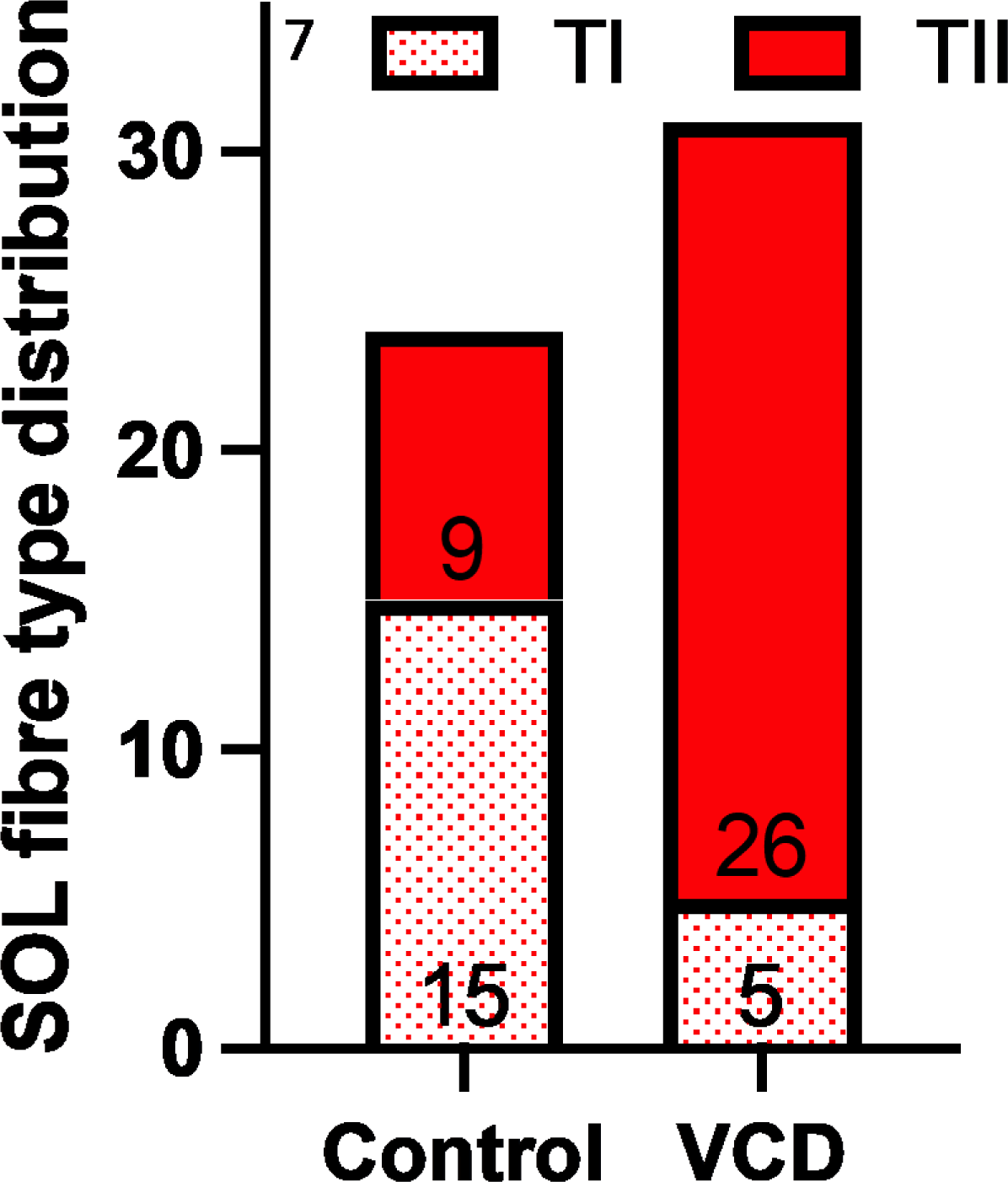
Fibre type distribution. The EDL for the VCD and control groups was comprised of 100% TII fibers. In the SOL muscle, the control group had ∼63-37% TI-TII split (TI, n=15; TII, n=9), while the VCD group had ∼16-84% TI-TII split (VCD TI, n=5; VCD TII, n=26), indicating a greater proportion of TII fibres in the VCD group.

### 4.4. The VCD group appears to have a greater proportion of TII fibers in the SOL

The EDL for the VCD and control groups was comprised of 100% TII fibers, typical of a mouse EDL[43]. In the SOL however, there was a shift to a greater proportion of TII fibres in the VCD group (Figure-7). This difference in fiber type distribution is likely driving the changes in contractile function observed in the SOL (and lack thereof in the EDL), indicating the SOL was faster to respond to the reduction in circulating 17β-estradiol. Greising et al., 2011 showed SOL normalized force was ∼30% lower with OVX, but the EDL was unaffected, they suggested the SOL might be slightly quicker to respond to 17β-estradiol manipulation than the EDL[10]. Additionally, Moran et al., 2007 reported no change in TI-TIIa-TIIx-TIIb fiber type distribution, however, once the TII fiber types are pooled together, there is a clear shift in TI-TII fiber type distributions, with 66% TII in the OVX group compared to sham, which mimics a similar trend to our current study[8]. Thus, in the above reported studies, as well as ours, 17β-estradiol deficiency has resulted in a shift toward a faster fiber-type distribution in the SOL.

### 4.5. Conclusion

With our VCD model of gradual ovarian failure, we characterized the effects of the perimenopausal transition on muscle contractility during a physiologically relevant phase of menopause. Our model approximates the natural transition to menopause in humans and highlights the physiological importance of time-course changes associated with menopause on neuromuscular function.

## Ethics statement

All procedures were approved by the Animal Care Committee of the University of Guelph.

## Data accessibility

Individual values of all supporting data are available upon request.

## Grants

This project was supported by the Natural Sciences and Engineering Research Council of Canada (NSERC) GAP and Heart and Stroke Foundation of Canada (HSFC) WGP.

## Author contributions

All authors contributed equally.

## Supporting information

chemical solutions

## Abbreviations

VCD: 4-Vinylcyclohexene diepoxide; a chemical for ototoxicity
SOL: Soleus muscle; a slower type skeletal muscle
EDL: Extensor digitorum longus muscle; a faster type skeletal muscle
CSA: Cross-sectional area
K_tr_: rate of force redevelopment; a measure of how rapid the muscle can generate force
K: instantaneous stiffness; a measure of the proportion of attached cross-bridges
L_0_: fiber length
SL: sarcomere length
OVX: ovariectomized; a model of completely removing the ovaries
Ca_2+_: Calcium

