## Supplementary material for "Effects of VCD-induced ovarian failure on single muscle fiber contractility in a mouse model of menopause": chemical solutions

The **dissecting solution** was composed of the following (in mM): K‐proprionate (250), Imidazole (40), EGTA (10), MgCl_2_·6H_2_O (4), Na_2_H_2_ATP (2), H_2_O.

The **storage solution** was composed of the following (in mM): K‐proprionate (250), Imidazole (40), EGTA (10), MgCl_2_·6H_2_O (4), Na_2_H_2_ATP (2), glycerol (50% of total volume after transfer to 50:50 dissecting:glycerol solution), as well as leupeptin (Sigma) protease inhibitors.

The **skinning solution** with Brij 58 was composed of the following (in mM): K‐proprionate (250), Imidazole (40), EGTA (10), MgCl_2_·6H_2_O (4), 1 g of Brij 58 (0.5% w/v).

The **relaxing solution** was composed of the following (in mM): Imidazole (59.4), K.MSA (86), Ca(MSA)2 (0.13), Mg(MSA)_2_ (10.8), K3EGTA (5.5), KH_2_PO_4_ (1), H_2_O, Leupeptin (0.05), Na_2_ATP (5.1), as well as leupeptin (Sigma) protease inhibitors.

The **pre‐activating solution** was composed of the following (in mM): KPr (185), MOPS (20), Mg(CH_3_COOH)_2_ (2.5), ATP (2.5). The activating solutions were composed of the same ingredients in various amounts depending on how much Ca^2+^ was needed in each solution. The composition of pCa 4.5 was as follows (mM):

The **activating solution** was composed of the following (in mM): Ca^2+^, Mg, EGTA (15), MOPS (80), ATP (5), CP (15), K (43.27), Na, and H_2_O. Ca^2+^, Mg, and Na concentrations were specific to the pCa level. These concentrations are listed below:

1. 4.5: Ca^2+^ (14.87), Mg (6.93), Na (13.23)
2. 5.5: Ca^2+^ (12.83), Mg (6.97), Na (13.4)
3. 5.7: Ca^2+^ (11.83), Mg (7), Na (13.4)
4. 6.2: Ca^2+^ (8.1), Mg (7.07), Na (13.4)
5. 6.4: Ca^2+^ (6.4), Mg (7.07), Na (13.4)
6. 6.6: Ca^2+^ (4.8), Mg (7.1), Na (13.4)
7. 7.0: Ca^2+^ (2.37), Mg (7.17), Na (13.4)

All solutions were adjusted to a pH of 7.0 with the appropriate acid (HCl) or base (KOH). The composition of solutions was determined by calculating the equilibrium concentration of ligands and ions based on published affinity constants (Fabiato and Fabiato, 1979). 250 units/ml of creatine phosphokinase was used in each activating solution.

**Homogenization buffer:** 61 mM tris (pH 6.8), 11% (*v/v*) glycerol, 2.78% (*w*/*v*) SDS, 5% 2-β-mercaptoethanol, and 0.02% (*w/v*) bromophenol blue.

**SDS-PAGE solutions**: The 7% separating gel consisted of 2 M tris HCL (pH 8.6), 50% glycerol, 10% sodium dodecyl sulfate (SDS), and 40% (*w/v*) acrylamide and N,N′ -methylenebis-acrylamide with a monomer to crosslinker ratio of 37.5:1. The stacking gel consisted of 500 mM tris HCL (pH 6.7), 10% SDS, and 40% (w/v) acrylamide and N,N′ -methylenebis-acrylamide with a monomer to crosslinker ratio of 37.5:1.
